# CDK12/13 inhibition disrupts nucleolar morphology and promotes aberrant expression of IGS transcripts

**DOI:** 10.1101/2025.03.01.640972

**Authors:** Rodolfo B. Serafim, Marina Volegova, Matthew Harlow, Ushashi Banerjee, Cibele Cardoso, Sanjukta Das, Bandana Sharma, Gabriel E. Zentner, Peter C. Scacheri, Tinghu Zhang, Nathanael S. Gray, Stephen Buratowski, Rani E. George

## Abstract

The transcription-associated cyclin dependent kinases (CDKs) 12 and 13 play important roles in the expression of RNAs involved in DNA damage repair and ribosomal proteins, yet their roles if any, in the regulation of ribosomal DNA expression are unclear. Here we show that these CDKs play critical roles in the processing and degradation of noncoding RNAs in the nucleolus and that inhibition of their function leads to the upregulation of unprocessed ribosomal RNA (rRNA) transcripts in the intergenic spacer (IGS) regions between ribosomal subunits. These upregulated RNAs are aberrantly polyadenylated by the TRAMP complex polymerase TENT4, and are represented as polyadenylated rings surrounding the nucleolus. The accumulation of these noncoding RNAs is associated with increased RNA polymerase II (Pol II)-mediated transcription coupled with failure of degradation by the MTREX (SKIV2L2) RNA helicase, a member of the RNA exosome complex whose expression is prematurely terminated by CDK12/13 inhibition. Altogether, these findings provide new insights into Pol II-mediated transcriptional regulation by CDK12/13 within the nucleolus.

## Introduction

The concept of transcriptional addiction has rendered RNA polymerase II (Pol II)-mediated gene expression an attractive therapeutic target in cancer^1,2^. Specifically, the cyclin-dependent kinases that facilitate various phases of the transcription cycle through phosphorylation of the C-terminal domain (CTD) of Pol II (transcription-associated CDKs or tCDKs) present potential targets whose activity can be disrupted by small molecule inhibition^3-8^. These include but are not limited to CDK7, responsible for transcription initiation, promoter clearance, and promoter-proximal pausing of Pol II, and CDK9 for Pol II pause release and elongation. CDK12 and CDK13, in association with partner cyclin K, facilitate the expression of full-length transcripts^9,10^. Apart from their roles in processivity and elongation rates of Pol II along gene bodies, these metazoan orthologs of the *S. cerevisiae*, Ctk1, recruit cleavage and polyadenylation factors and prevent read-through transcription and abnormal 3’-end processing^5,9-12^. Unlike CDK7 and CDK9 that affect global transcription, CDK12 specifically regulates the expression of genes involved in DNA damage repair while CDK13 is involved in small nuclear and small nucleolar RNA regulation, enabling nuclear RNA surveillance^7,11-13^. We and others have characterized the on-target effects of selective covalent inhibitors of tCDKs and established their tumor-inhibiting capabilities^7,14,15^. We identified CDK12 as a therapeutic vulnerability in neuroblastoma and osteosarcoma^7,15^ and demonstrated that the CDK12/13 inhibitor THZ531 (covalently targeting CDK7, 8.5 µM; CDK12, 0.158 µM; CDK13, 0.069 µM)^16^ and the more selective E9 (targeting CDK9 through ATP competition and CDK12 through covalent binding)^17^ abrogated metastatic outgrowth of osteosarcoma cells^15^. Following THZ531 treatment, Pol II phosphorylation was decreased and transcription elongation and termination disrupted in a gene length-dependent manner with long (>45 kb) genes undergoing early termination through the increased use of intronic polyadenylation sites, premature cleavage and polyadenylation, while short genes retain 3’-downstream readthrough^7^. CDK12 activity also controls the translation and phosphorylation of proteins involved in ribosome biogenesis and ribosomal RNA (rRNA) processing respectively^7,18^. While the effects of CDK12/13 inhibitors on the protein coding genome have been studied extensively, their consequences on ribosomal DNA expression are not well understood. Here, we identify the transcriptional changes which occur in the nucleolus as a result of small molecule inhibitors that perturb Pol II function through disrupting CDKs 12/13. We demonstrate that cancer cells express low levels of non-polyadenylated intergenic rRNAs at baseline and that in response to CDK12/13 inhibition, these transcripts are not only upregulated, but are also aberrantly polyadenylated, leading to altered nucleolar structure and function. Finally, we determine that the aberrant transcript accumulation is due in part to increased Pol II activity coupled with the loss of RNA exosome activity, thereby preventing degradation.

## Results

### CDK12/13 inhibition disrupts nucleolar morphology and rRNA processing

To understand the roles of CDK12/13 on rDNA transcription, we first analyzed the impact of CDK12/13 inhibition on nucleolar structure using sub-lethal concentrations of the CDK12/13 inhibitor, E9 on human osteosarcoma cells. The nucleolus is a membrane-less organelle formed through liquid-liquid phase separation and is comprised of 3 different functional layers - the fibrillar center (FC) containing rDNA and responsible for rRNA transcription, the dense fibrillar center (DFC) for rRNA maturation and the granular center (GC) for ribosomal assembly (**Figure 1A**). The three layers are surrounded by a perinucleolar heterochromatin ring marked by histone 3 lysine 9 trimethylation (H3K9me3) that physically separates the nucleolus from the rest of the nucleus, thereby preventing interference from other cellular processes that might disrupt ribosome assembly^19-21^. We analyzed the expression of surrogate architectural markers of each of the nucleolar layers by immunofluorescence and noted that E9 treatment led to nucleolar contraction with aberrant localization of the sub-compartmental proteins; NOLC1 and NPM1 staining marking the FC and GC respectively, were decreased overall with partial dissolution into the nucleoplasm and localization to the peri-nucleolar space (**Figure 1A**). The DFC, where most rRNA maturation takes place, appeared to be most affected, with decreased expression and dispersal of FBL through the nucleus. In addition, decreased staining of H3K9me3 at the nucleolar periphery was observed, suggesting disruption of the heterochromatin barrier (**Figure 1B**). The alterations in nucleolar morphology implied that rRNA transcription may be affected and hence, we next examined the effects of CDK12/13 inhibition on rRNA expression. Using rRNA in situ hybridization (RNA-ISH), we noted that following exposure to E9, 28S and 18S rRNAs appeared to be much more punctate compared to their more dispersed signal in DMSO-treated cells, indicating nucleolar contraction (**Figure 1C**, **Supplementary** Figure 1A). However, quantification of rRNA expression by reverse transcription-quantitative polymerase chain reaction (RT-qPCR) showed that E9 treatment caused no significant change in the expression of these genes (**Figure 1D**). CDK12 degradation using a selective degrader, BSJ-4-116, derived from the covalent CDK12 inhibitor, THZ531^16^ also did not significantly affect their expression (**Figure 1D**). These results suggest that factors other than rDNA expression may account for the aberrant nucleolar morphology observed with CDK12/13 inhibition.

**Figure 1.**
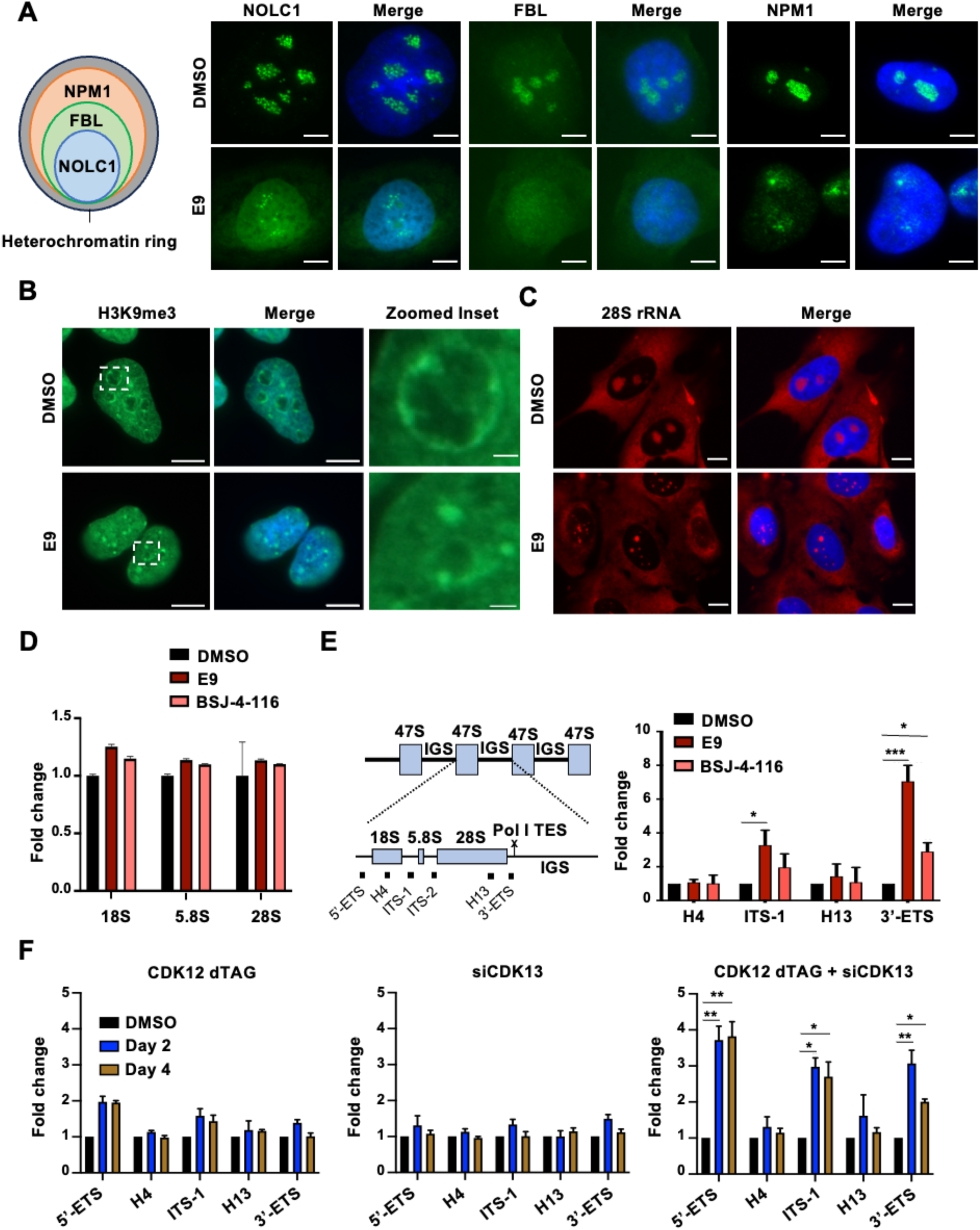
CDK12/13 inhibition disrupts nucleolar morphology. (A) Schematic model of nucleolar structure (left). Immunofluorescence (IF) analysis of the indicated nucleolar marker expression in OS cells incubated with E9 (200 nM x 6 h). Scale bar, 5 μm. (B) IF analysis of H3K9me3 staining in OS cells treated as in (A). Scale bar, 10 μm, inset, 1 μm. (C) RNA-ISH analysis of 28S rRNA in OS cells treated as in (A). Scale bar, 10 μm. (D) RT-qPCR analysis of rRNA expression in OS cells treated with E9 as in A or BSJ-4-116 (500 mM x 6 h). GAPDH was used as loading control. Data represent mean ± standard deviation (S.D.) of 3 independent replicates. (E). RT-qPCR analysis of rRNA expression (right) at the regions corresponding to the schematic model of the rDNA tandem repetitive structure (left) in OS cells treated as in (D). Blue boxes represent the 47S rDNA region. H4 and H13 represent primers binding to 18S and 28S, respectively. ITS-1 (Intragenic Spacer 1); ITS-2 (Intragenic Spacer 2); 3’-ETS (3’-Externally Transcribed Spacer); IGS (intergenic spacer); Pol I TES (Pol I Transcription End Site). GAPDH was used as loading control. Data represent mean ± S.D. of 3 independent replicates. \**P* < 0.05, and \*\*\**P* < 0.0001, one-way ANOVA followed by Dunnett’s multiple comparisons test. (F) RT-qPCR analysis of rRNA expression in 143B CDK12 dTAG cells transfected with siControl and treated with DMSO or 1 µM dTAG13 (left), transfected with siControl or siCDK13 (middle), and transfected with siCDK13 and treated with DMSO or 1 µM dTAG13 (right). dTAG13 treatment and CDK13 knockdown were performed for 2 or 4 days. GAPDH was used as loading control. Data represent mean ± S.D. of 2 independent replicates. \**P* < 0.05, and \*\**P* < 0.001, one-way ANOVA followed by Dunnett’s multiple comparisons test.

To explain the abnormalities in nucleolar architecture observed with CDK12/13 depletion, we analyzed transcription along the entire rRNA sequence. Human ribosomal DNA is made up of approximately 400 repeats, each 45 kilobase (kb) repeat comprising the 18S, 5.8S and 28S rRNA genes (proximal 13 kb) separated by two intragenic spacers (ITS-1, between 18S and 5.8S and ITS-2, between 5.8S and 28S), and flanked by 5′ and 3′ external transcribed spacers (ETS), which are removed during processing. The rRNA repeats are separated by intergenic spacers (IGS) (30 kb) that contain repetitive sequences with enhancer and promoter elements that contribute to the maintenance of nucleolar structure and organization^22-25^ (**Figure 1E**). Interestingly, we observed rRNA transcript accumulation at the ITS-1 and 3’-ETS regions with both E9 and BSJ-4-116 (**Figure 1E**). To confirm that these effects were indeed the result of on-target kinase inhibition, we assessed the effects in osteosarcoma cells in which we introduced a degradation tag (dTAG)^26^ to rapidly deplete CDK12, as well as a short-interfering RNA (siRNA) directed against CDK 13 (**Supplementary** Figures 1B-C). While downregulation of CDK12 and to a lesser extent, CDK13 individually led to minor effects, the changes in rRNA expression were only evident after depletion of both kinases (**Figure 1F**), suggesting that the activity of both kinases must be inhibited to effectively disrupt rRNA processing. Altogether, these observations underscore the critical role of CDK12/13 activity in preserving nucleolar structure and ensuring proper rRNA processing.

### CDK12/13 inhibition leads to aberrant rRNA expression at the H16 IGS locus

To delineate the extent of the aberrantly increased transcripts following CDK12/13 inhibition, we systematically analyzed RNA expression across the entire 30 kb IGS region in cells treated with E9 (**Figure 2A**). Of note, the duration of treatment was designed to observe the immediate on-target effects of CDK12/13 inhibition (loss of Pol II phosphorylation) but preceded the deleterious downstream effects of drug treatment, such as cell death, which was evident following 24-hour washout of the drug (**Supplementary** Figure 2). We observed a significant increase in the total expression of transcripts generated from the IGS, with the most pronounced upregulation observed at the H16 locus (**Figure 2A**). The increase at H16 occurred in a time-dependent manner in response to drug treatment (**Figure 2B**). The effects at H16 were also noticeable following treatment with the CDK12 degrader, BSJ-4-116 and CR8, a degrader of the cyclin K partner of CDK12^27,28^ (**Figure 2C**), supporting our observation that CDK12/13 inhibition increases the expression of aberrant ribosomal IGS transcripts.

**Figure 2.**
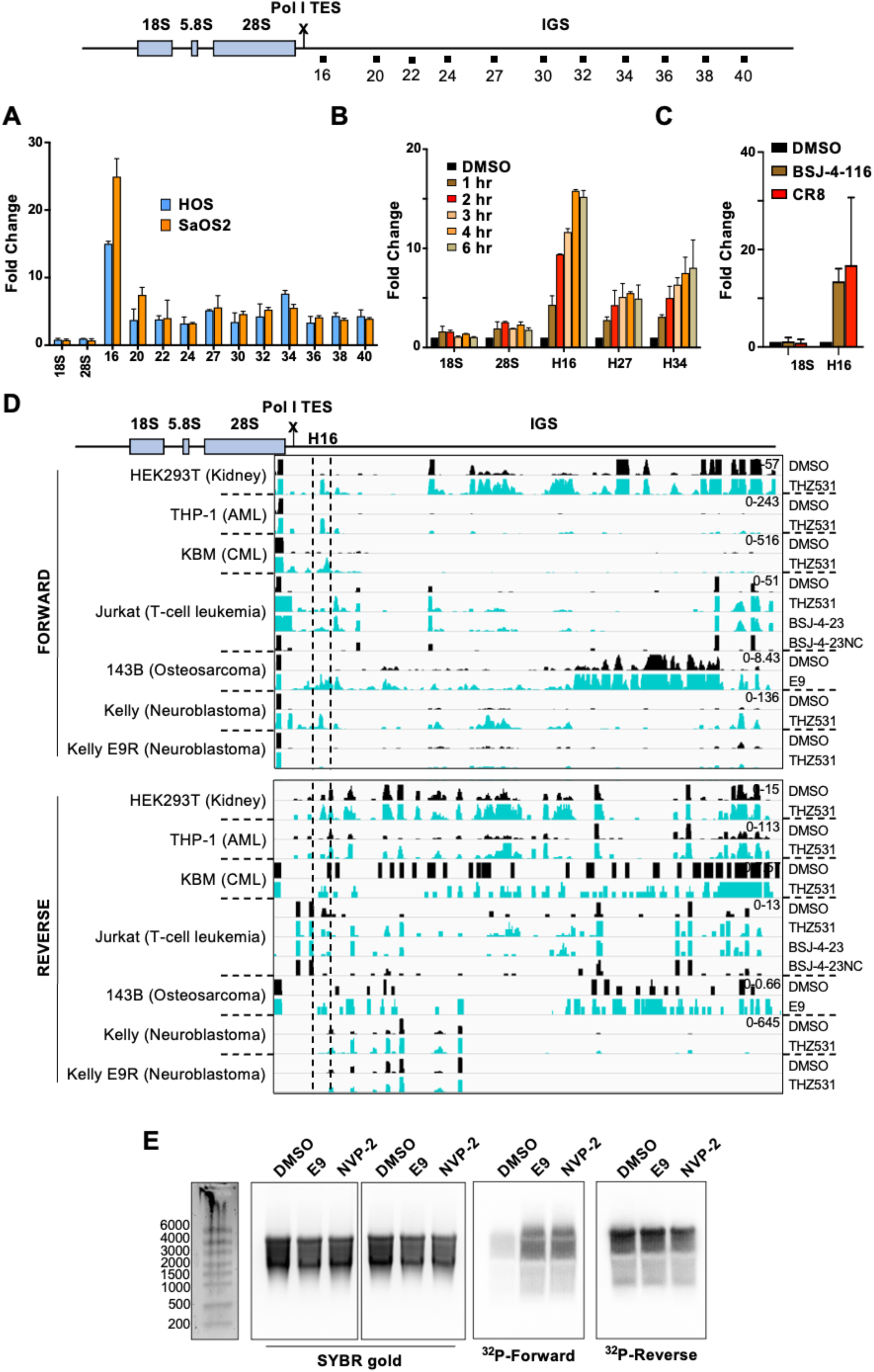
CDK12/13 inhibition results in increased IGS transcript expression. (A) RT-qPCR analysis of 18S, 28S and IGS expression (at the regions indicated in the cartoon above) in OS cells treated with E9 (200 nM x 6 h). (B) RT-qPCR analysis of the indicated rRNAs in OS cells treated with E9 (200 nM) for the indicated times. Data represent mean ± S.D. of 2 independent replicates in A and B. (C) RT-qPCR analysis of 18S and H16 IGS region in OS cells incubated with BSJ-4-116 (500 nM x 6 h) or CR8 (600 nM x 6 h). Data represent mean ± S.D. of 3 independent replicates. GAPDH was used as loading control in all three assays. (D) Integrative Genomics Viewer (IGV) tracks of RNA expression in forward and reverse orientations from analysis of published RNA sequencing data sets of the indicated cell lines treated with CDK12/13 inhibitors at the following doses and durations: HEK293T: THZ531 (400 nM x 6 h); THP-1: THZ531 (200 nM x 6 h); KBM7: THZ531 (600 nM x 5 h); JURKAT: THZ531 (250 nM x 8 h); 143B: E9 (200 nM x 6 h); KELLY: 400 nM (THZ531 x 6 h). The y-axis scale indicates the range of normalized read coverage for each sample. Data set accession numbers are given in Supplemental Table 2. Dashed lines, H16 locus. (E) Northern blot analysis of RNA from OS cells treated with E9 (200 nM x 6 h) or NVP-2 (50 nM x 6 h). Equal loading across the samples is indicated by SYBR gold staining (left panels) of total RNA. The H16 transcripts were detected using radiolabeled probes targeting the forward (^32^P-Forward) or reverse (^32^P-Reverse) strands.

To determine whether the effects of CDK12/13 inhibition on IGS transcription extended beyond osteosarcoma cells, we analyzed publicly available bi-directional RNA sequencing data from transformed and cancer cell lines exposed to various inhibitors that targeted these kinases^7,15,27,29,30^. Upon treatment with either CDK12 kinase inhibitors (THZ531 or E9) or a selective degrader, BSJ-4-23^27^, significantly increased expression of transcripts emanating from the H16 locus was seen in multiple cell lines, irrespective of cell lineage or tumor origin (**Figure 2D**). However, H16 upregulation was not observed in E9-resistant neuroblastoma cells (E9R) that express the CDK12 C1039 mutation at the covalent binding site of E9^7^, indicating the on-target effect of CDK12 inhibitors. Additionally, the IGS transcriptional profile of Jurkat leukemia cells treated with the negative control CDK12 degrader (BSJ-4-23-NC^27^) phenocopied the solvent condition, demonstrating no effect on H16 expression (**Figure 2D**).

CDK12/13 inhibition resulted in consistent upregulation of transcription at the H16 locus in the forward rDNA strand, although the same finding was not observed in reverse strand sequences (**Figure 2D**). To confirm the direction of the H16 transcript, we performed northern blot analysis using locus-specific forward and reverse oligonucleotide probes (**Figure 2E**). Binding of the reverse probe to RNA in DMSO-treated cells suggested the presence of an anti-sense or reverse transcript at this locus, consistent with the anti-sense IGS transcription previously described by Abraham *et al^31^* which was unchanged following E9 treatment. On the other hand, while there was no binding of the forward probe under DMSO conditions, a transcript was seen following E9 treatment, confirming the presence of the aberrant H16 transcript and its forward orientation. A similar result was obtained in cells treated with the CDK9 inhibitor NVP-2^32^ in keeping with the reported role of CDK9 in the nucleolus^33^. Thus, CDK12/13 inhibition leads to aberrant upregulation of a sense transcript at the H16 locus irrespective of cell lineage.

### The aberrantly expressed IGS rRNAs are transcribed by Pol II

rRNA genes are transcribed by Pol I, while RNA expression at the IGS is thought to be mediated by Pol II^31^. Pol II-dependent noncoding RNAs (ncRNAs) are transcribed in the reverse direction and prevent Pol I from generating forward intergenic ncRNAs that disrupt nucleolar organization and rRNA expression. Given that the H16 IGS RNA was transcribed in the forward direction, it was unclear whether it was due to run-on Pol I activity or nucleolar Pol II activity. We reasoned that if Pol I was responsible for the upregulated H16 expression, inhibition of Pol I with low dose actinomycin (LAD) or the Pol I inhibitor CX-5461 in E9-treated cells would decrease H16 expression. However, H16 downregulation was not observed with either of these compounds suggesting the lack of involvement of Pol I in generating the IGS transcripts (**Figure 3A**). In addition, we treated the cells with the pan-CDK inhibitor flavopiridol which induces complete blockade of Pol II activity by targeting multiple cell cycle and transcriptional CDKs (CDKs 1, 2, 4, 6, 7, 9)^34^. As with E9, increased RNA expression at H16 was also seen with flavopiridol, however, unlike E9-treated cells, H16 overexpression was rescued by LAD and CX-5361, implying that these transcripts arose from Pol I-mediated IGS readthrough (**Figure 3A**).

**Figure 3.**
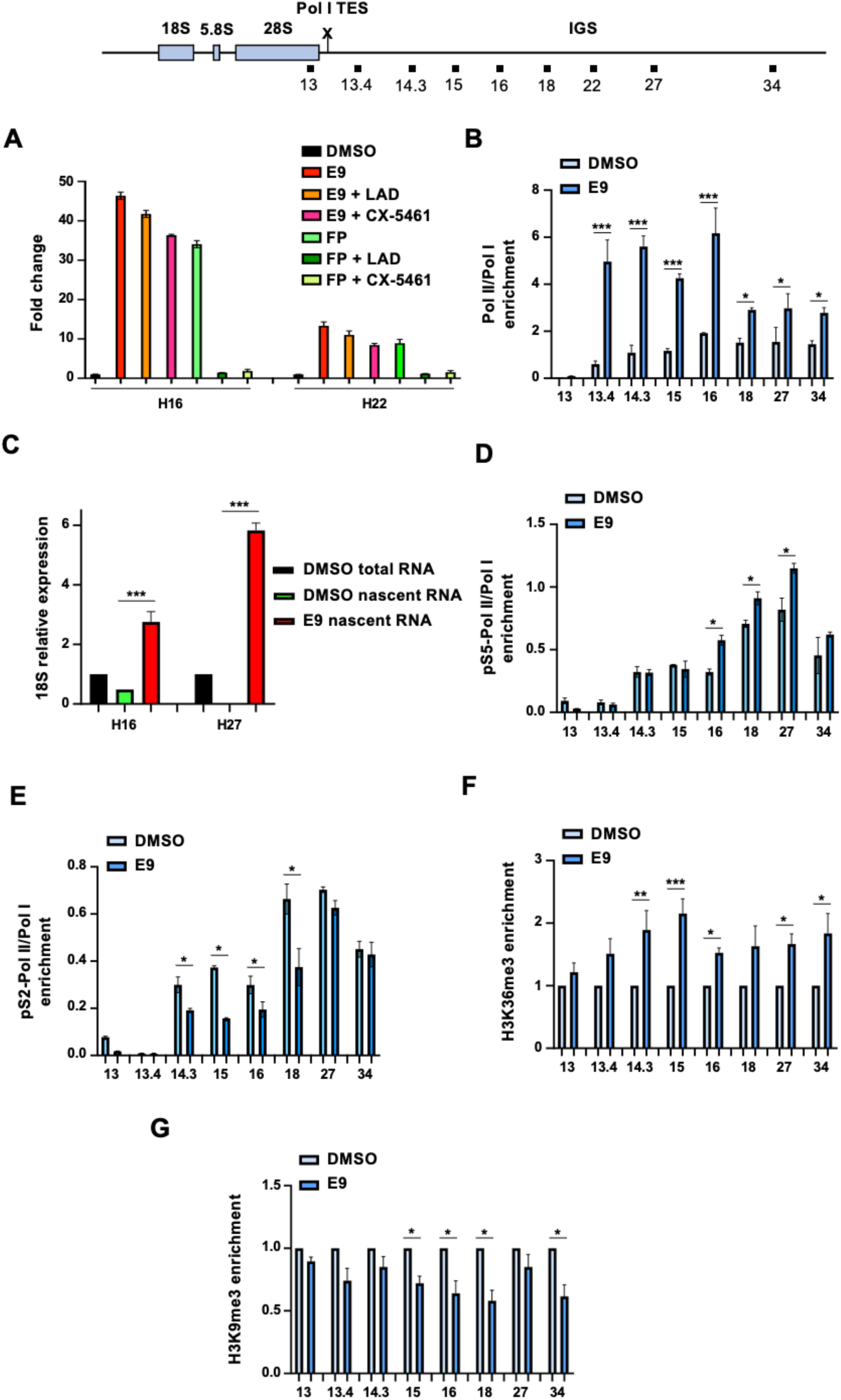
IGS RNAs are transcribed by Pol II. (A) RT-qPCR analysis of H16 and H22 rRNA expression in HOS cells treated for 6 h with flavopiridol (FP) (2 µM), actinomycin D (LAD) (50 ng/ml), CX-5461 (200 nM) and/or E9 (200 nM). GAPDH was used as loading control. Data represent mean ± S.D. of 2 independent replicates. (B) ChIP-qPCR analysis of Pol II and Pol I occupancies at the 28S (H13) and IGS regions (at the regions indicated in the cartoon above) in HOS cells treated with E9 (200 nM x 6 h). The ratio of Pol II to Pol I binding was calculated as (% of input for Pol-II immunoprecipitation)/(% of input for Pol-I immunoprecipitation)/(% of input for mock IgG immunoprecipitation). Data represent mean ± S.D. of 2 independent replicates. \**P* < 0.05, and \*\*\**P* < 0.0001, Student’s t test. (C) Nascent RNA emanating from H16 and H22 regions in HOS cells treated with E9 (200 nM x 6 h). 18S was used as loading control. Data represent mean ± S.D. of 2 independent replicates. \*\*\**P* < 0.0001, one-way ANOVA followed by Dunnett’s multiple comparisons test. (D-E) ChIP-qPCR analysis of pS5-Pol II (D) or pS2-Pol II (E) occupancies at the 28S and IGS regions in HOS cells treated as in (B). pS5- and pS2-Pol-II/Pol-I ratio calculated as in (B). Data represent mean ± S.D. of 2 independent replicates. \**P* < 0.05, Student’s t test. (F-G) ChIP-qPCR analysis of H3K36me3 (F) or H3K9me3 (G) occupancies at the 28S and IGS regions in HOS cells treated as in (B). Data represent mean ± S.D. of 2 independent replicates. \**P* < 0.05, \*\**P* < 0.001, and \*\*\**P* < 0.0001, Student’s t test.

To further corroborate these results, we analyzed the relative occupancies of each RNA polymerase along the rDNA locus through chromatin immunoprecipitation and quantitative polymerase chain reaction (ChIP-qPCR) analysis. As expected, the Pol II to Pol I occupancy ratio was ≤1 at the rRNA genes, ITS-1 and 3’-ETS regions under control conditions, indicating preponderance of Pol I at these regions, which remained unchanged following E9 treatment (**Figure 3B**, **Supplementary** Figure 3A). On the other hand, Pol II occupancy was predominant at the IGS as seen by the Pol II/Pol 1 ratio; moreover, it was markedly increased following CDK12/13 inhibition (**Figure 3B**). The increased Pol II binding arose downstream of the rRNA transcription termination site and extended throughout the IGS, but with the most pronounced effects seen just after H13 to H16 (**Figure 3B**; **Supplementary** Figure 3A). The observed increase in Pol II binding was reflected in increased nascent RNA levels at the H16 locus in E9-treated cells (**Figure 3C**), confirming the upregulated Pol II transcriptional activity following CDK12/13 inhibition. Both active forms of Pol II, the initiation-associated phosphorylated serine 5 (pS5), and the elongation-associated phosphorylated serine 2 (pS2), were significantly enriched in the IGS compared to the 28S gene in control conditions (**Figure 3D-E**). Treatment with E9 resulted in increased levels of pS5-Pol II occupancy in the region comprising H16 to H27 (**Figure 3D**), with a reduction in pS2-Pol II occupancy in H14 to H18 regions (**Figure 3E**), together pointing to an elongation defect following CDK 12/13 inhibition. In addition, we observed concomitant increased enrichment of H3K36me3, a chromatin mark coupled to active transcription, but not thought to be associated with Pol I activity^35-37^ (**Figure 3F**). Decreased occupancy of the H3K9me3, a marker of repressed chromatin^19^, was also seen following E9 treatment (**Figure 3G**), confirming the results of our prior immunofluorescence assay (**Figure 1B**). Importantly, these findings were consistent across multiple cell lines and following treatment with either E9 or THZ531 (**Supplementary** Figure 3B-D). Altogether, findings attest to the predominance of Pol II-mediated transcription along the IGS and support a model whereby small molecule CDK12/13 inhibition induces a more transcriptionally permissive chromatin environment leading to the aberrant expression of ncRNAs which could ultimately disrupt nucleolar organization.

### IGS transcripts are aberrantly polyadenylated following CDK12/13 inhibition

ncRNAs undergo polyadenylation to increase transcript stability, modulate interactions with other proteins or RNA molecules, and regulate subcellular localization^38,39^. To determine if the IGS transcripts were polyadenylated, we examined the change in polyadenylated (poly(A)) RNA expression across the rDNA locus following E9 treatment. We observed an increase in poly(A) RNA expression in the IGS that phenocopied the change in total RNA expression following drug treatment, with the highest levels occurring in the H16 region. (**Figures 4A and 2A**). We next mapped location and length of the polyadenylated tail of the H16 transcript using a reverse primer specific to this region. (**Figure 4B**). Prior to E9 treatment, we observed a minimal amount of polyadenylated transcript in the forward orientation at the H16 locus, consistent with low-level pervasive transcription (**Figure 4B**). However, following E9 treatment, there was a significant increase in polyadenylated H16 transcript expression with a poly(A) tail length variance of 70-100 nucleotides, well within the range of other polyadenylated RNAs produced by Pol II (**Figure 4B**)^40,41^. Following sequencing and subsequent validation by 3’-RACE (Rapid Amplification of cDNA Ends), we further defined the polyadenylation site of the H16 transcript and localized its termination site to approximately 2.7 kilobases downstream of the Pol I termination site (**Figure 4C**). Selective depletion using siRNAs directed to the H16 locus in E9-treated cells led to a reduction in expression at this region, while H22 to H42 locus expression was not affected, suggesting that multiple aberrant transcripts rather than a single transcript are produced along the IGS following CDK12/13 inhibition and may contribute to the nucleolar dysfunction (**Figure 4D**).

**Figure 4.**
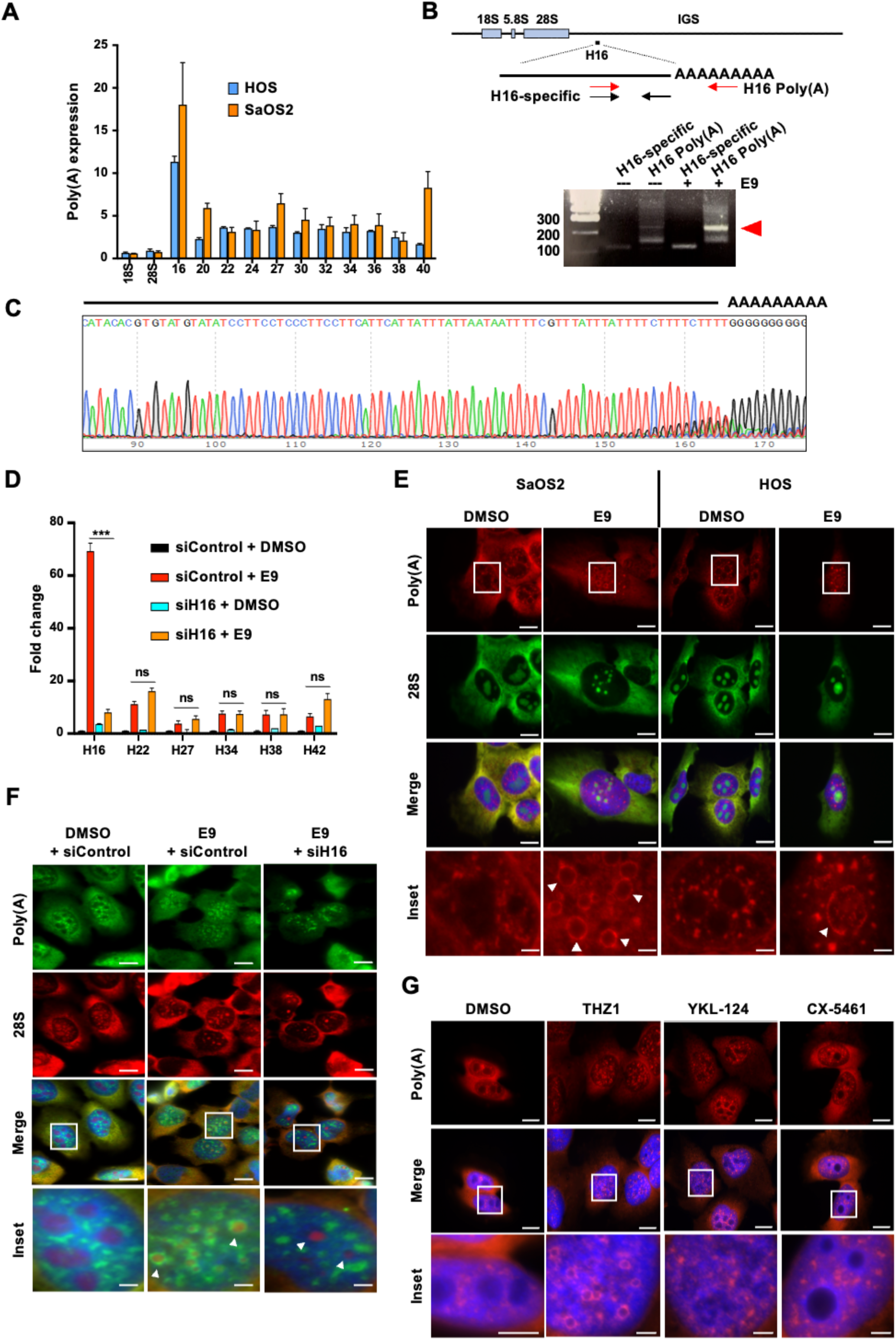
IGS transcripts are polyadenylated following CDK12/13 inhibition. (A) RT-qPCR analysis using IGS-forward specific primers and a universal reverse primer to detect the poly(A) tail in OS cells treated with E9 (200 nM x 6 h). Data represent mean ± S.D. of 2 independent replicates. (B) PCR analysis using H16-specific primers and poly(A) primers in OS cells treated as in (A). The arrowhead indicates the polyadenylated H16 RNA. (C) Sanger sequencing analysis of DNA samples purified from (B). (D) RT-qPCR analysis of IGS region in OS cells transfected with siRNA against H16 RNA and treated with E9 (200 nm x 6 h). Data represent mean ± S.D. of 3 independent replicates. \*\*\**P* < 0.0001, one-way ANOVA followed by Dunnett’s multiple comparisons test. ns, not significant. (E) IF analysis of poly(A) and 28S rRNA in OS cells incubated with E9 (200 nM x 6 h). Arrow heads show pAR rings. Scale bar, 10 μm, inset, 2 μm. (F) IF analysis of same markers as in (E) in SaOS2 cells transfected and treated as in (D). Arrow heads show pAR rings or its absence in H16-knockdown cells. Scale bar, 10 μm, inset, 2 μm. (G) IF analysis of poly(A) in OS cells incubated with THZ1 (200 nM x 6 h), YKL-124 (200 nM x 6 h), or CX-5461 (200 nM x 6 h). Scale bar, 10 μm, inset, 2 μm.

We next determined the cellular localization of the H16 transcript using RNA ISH with a probe complementary to the poly(A) tail. Under normal conditions, we observed punctate foci throughout the nucleoplasm and a more dispersed signal in the cytoplasm that co-localized with 28S rRNA (**Figure 4E**). Importantly, the poly(A) signal was excluded from the nucleolus. Following E9 treatment, there was a dramatic change in poly(A) RNA expression with decreased cytoplasmic distribution, larger nucleoplasmic foci, and the appearance of distinct rings surrounding the nucleolus (**Figure 4E**). Similar to the kinetics of IGS transcript accumulation, the polyadenylated RNA (pAR) ring phenotype appeared in a concentration- and time-dependent manner (**Supplementary** Figure 4A-B). Consistent with the rescue of the aberrantly increased H16 expression, H16 siRNA knockdown resulted in loss of the pAR ring phenotype in E9-treated cells (**Figure 4F**), suggesting that the pAR rings represented the aberrantly transcribed polyadenylated IGS RNAs. However, the 28S signal remained punctate, indicating persistent nucleolar dysfunction, as confirmed by NPM1 staining (**Supplementary** Figure 4C). Treatment with THZ1 also resulted in a similar polyadenylated ring phenotype, in keeping with its known activity against CDKs 12 and 13^42^, while a selective CDK7 inhibitor, YKL-124^42^ or the Pol I inhibitor, CX-5461^43,44^, did not induce the perinucleolar ring structure (**Figure 4G**). Together, these results suggest that the pAR rings are an on-target effect of CDK12/13 inhibition and represent the polyadenylated aberrant transcript at the H16 locus.

### RNA exosome dysfunction contributes to aberrant IGS transcript accumulation

Transcript expression is influenced by both RNA synthesis and decay^45^. Under normal conditions, IGS transcripts are nascently transcribed at low levels and their lack of stable expression suggests that they are rapidly degraded by the RNA exosome, similar to other pervasively transcribed RNAs^46,47^. We therefore hypothesized that, in addition to increased RNA synthesis, perturbations in RNA degradation may also contribute to the upregulation of IGS transcripts following CDK12/13 inhibition. Nucleolar RNA degradation is carried out mainly by the Trf4/Air2/Mtr4p polyadenylation (TRAMP) complex, that detects aberrantly transcribed or mis-processed rRNAs and processes them for exosomal degradation^48-50^. The nucleoplasmic NEXT (Nuclear Exosome Targeting) and the PAXT (Polyadenylated Exosome Targeting) processing complexes degrade non-polyadenylated Pol II transcripts (i.e. upstream anti-sense promoter transcripts) and pre-mRNAs^51^, respectively. All three complexes sense their substrates through the action of RNA-binding proteins and are ultimately bound by the RNA helicase MTREX (SKIV2L2) which eliminates secondary structure and transmits the RNAs into the central channel of the RNA exosome^50^.

To determine the consequences of CDK12/13 inhibition on RNA exosome function, we first analyzed the expression of genes encoding the nuclear RNA exosome co-factor complexes following CDK12/13 inhibition in OS and neuroblastoma cell lines from publicly available gene expression data (GSE113314^7^ and GSE132233^15^). As with most protein-coding genes, the expression of RNA exosome and co-factor genes was suppressed with drug treatment (**Figure 5A**). Because CDK12/13 inhibition affects gene expression in a length-dependent manner, preferentially targeting long transcripts^7,52^, and the TRAMP complex genes are significantly longer than those encoding subunits of the other processing complexes (**Figure 5B**), we reasoned that loss of TRAMP complex gene expression could be due to premature termination. Analysis of 3′-poly(A) RNA-sequencing data from neuroblastoma cells exposed to THZ531^15^ indeed revealed that the genes encoding TRAMP complex components ZCCHC7, ZCCHC714 and MTREX were prematurely terminated at canonical intronic polyadenylation sequences (**Supplementary** Figure 5A).

**Figure 5.**
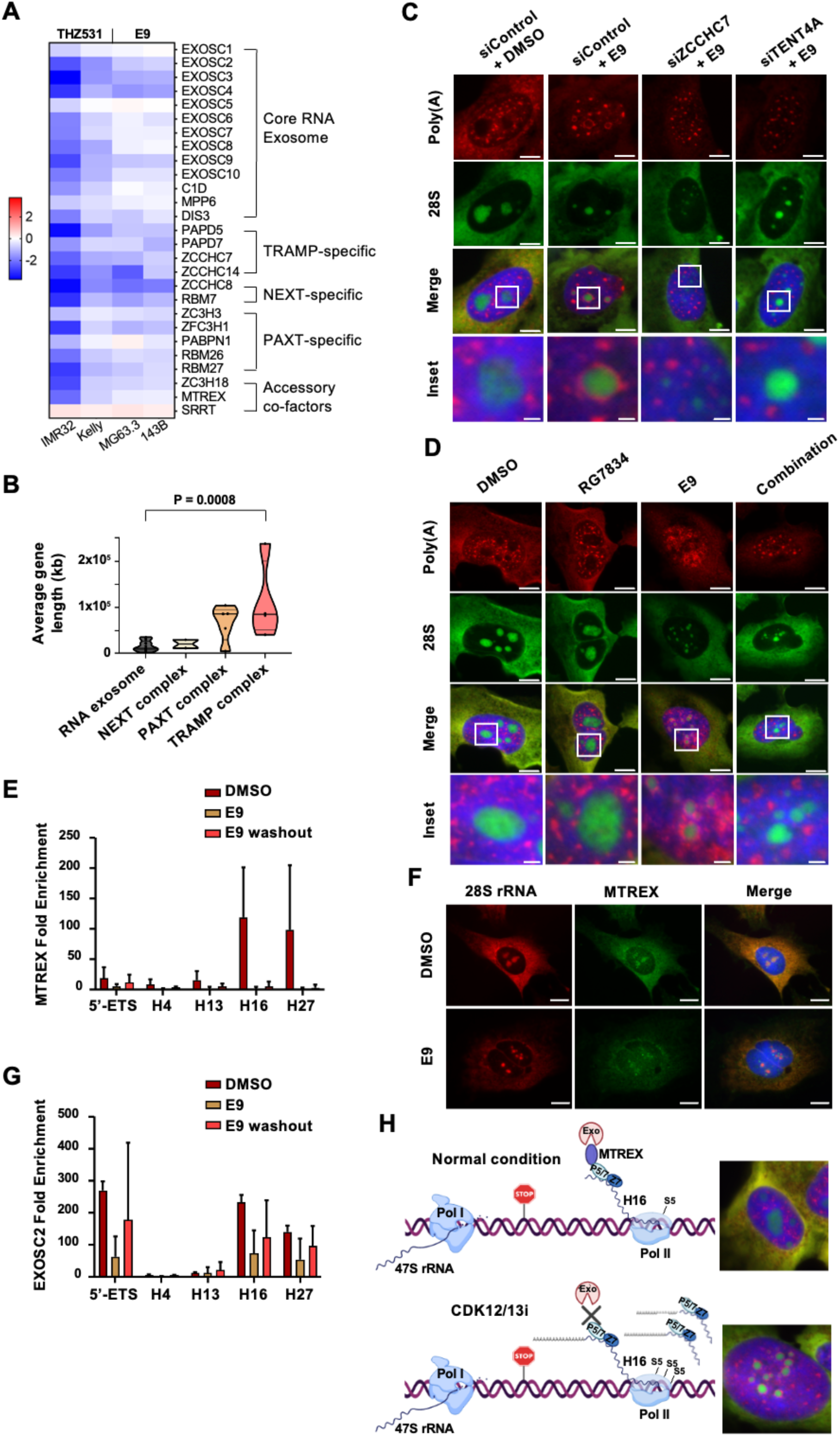
IGS transcripts accumulate following CDK12/13 inhibition due to RNA exosome dysfunction. (A) Heatmap showing the differential expression levels of RNA exosome and nuclear exosome co-factors in IMR32, Kelly, MG63.3, and 143B cells treated with THZ531 (500 nM x 6 h) or E9 (200 nM x 6 h) compared to DMSO treated control. Expression levels were obtained from public RNA-seq datasets GSE113314^7^ and GSE132233^15^ from the GEO database. Differential expression is represented as log2(fold change), with upregulation shown in red and downregulation in blue. (B) Violin plot showing the average gene length of core RNA exosome, NEXT, PAXT and TRAMP components. (C) IF analysis of poly(A) and 28S rRNA in OS cells transfected with siRNA against ZCCHC7 or TENT4A mRNA and incubated with E9 (200 nM x 6 h). Scale bar, 5 μm, inset, 1 μm. (D) IF analysis of the same markers as in (C) in OS cells treated with E9 (200 nM x 6 h) and/or RG-7834 (10 μM x 6 h). Scale bar, 5 μm, inset, 1 μm. (E) RIP-qPCR analysis of MTREX binding to 18S, 28S, H16 and H27 rRNAs in OS cells treated with E9 (200 nM x 6 h). Data represent mean ± S.D. of 2 independent replicates. RNA enrichment was calculated as (% of input of rRNA or IGS)/(% of input of GAPDH). (F) IF analysis of 28S rRNA and MTREX in OS cells treated as in (E). Scale bar, 5 μm. (G) RIP-qPCR analysis of EXOSC2 binding to 18S, 28S, H16 and H27 rRNAs in OS cells treated as in (E). Data represent mean ± S.D. of 2 independent replicates. RNA enrichment was calculated as in (E). (H) Schematic model for pAR rings accumulation following CDK12/13 inhibition in OS cells.

In addition to the RNA helicase MTREX, and the RNA-binding proteins ZCCHC7 or ZCCHC14, the non-canonical polymerases TENT4A (PAPD5) or TENT4B, also make up the TRAMP complex. Aberrantly produced nucleolar RNAs are bound by ZCCHC7 or ZCCHC14, which are then polyadenylated by TENT4A and directed to the exosome by MTREX. The TRAMP complex must retain both RNA binding and polymerase activities to target substrate RNAs to the RNA exosome^50^. The polyadenylation of the IGS transcripts suggested that although TRAMP complex components were downregulated with CDK12/13 inhibition, processes mediated by ZCCHC7/ZCCHC14 and TENT4A – RNA binding and polyadenylation respectively - remained functional. As such, we reasoned the pAR phenotype would be eliminated following loss of either activity. In E9-treated cells, either ZCCH7 or TENT4A siRNA knockdown was sufficient to completely reverse the pAR phenotype (**Figure 5C**, **Supplementary** Figure 5B). Treatment with small-molecule TENT4A/B inhibitor, RG-7834^53^ also phenocopied the effect of TENT4A knockdown, resulting in the elimination of pAR rings (**Figure 5D**). Importantly, ZCCH7 or TENT4A depletion alone did not affect rRNA expression or cell viability (**Supplementary Figure 5B-D**) suggesting that the elimination of pAR rings was not due to generalized toxicity.

Thus far, our data suggest that the effect of CDK12/13 inhibition on TRAMP complex function occurs downstream of TENT4A/B-dependent polyadenylation, at the MTREX helicase step. We therefore investigated the ability of MTREX to bind to and transmit transcripts to the exosome using RNA immunoprecipitation followed by qPCR (RIP-qPCR). Under solvent conditions, MTREX was robustly associated with transcripts originating from the H16 region of the IGS, suggesting that MTREX facilitates immediate destruction of these pervasive Pol II transcripts (**Figure 5E**). However, following treatment with E9, RNA binding of MTREX in the nucleolus was eliminated (**Figure 5E**). This effect persisted even upon drug washout, arguing that the RNA binding defect is a direct consequence of covalent inhibition of CDK12/13 by E9. In agreement with this result, the MTREX protein was excluded from the nucleolar compartment following treatment (**Figure 5F**). Finally, we noted that the ability of the RNA exosome (EXOSC2) to bind polyadenylated IGS transcripts was reduced following continuous E9 treatment, implying that the upstream function of MTREX had been compromised (**Figure 5G**). Overall, these data suggest that the accumulation of aberrantly polyadenylated IGS transcripts following small-molecule CDK12/13 inhibition is partially due to the inability of the MTREX helicase to process and transmit TENT4-polyadenylated RNAs to the RNA exosome (**Figure 5H**).

## Discussion

Transcriptional CDK inhibition has been established as a promising therapeutic modality against cancer, yet its clinical translation has been hampered by generalized and often unexplained toxicity. Here, we sought to characterize the consequences of transcriptional CDK inhibition on nucleolar structure and function. We demonstrate that CDK12 and CDK13, key CDKs involved in Pol II-mediated transcription, act in concert to regulate the expression of aberrant ncRNAs in the nucleolus. CDK12/13 inhibition leads to increased Pol II occupancy along the IGS region, with the resultant overexpression of a discrete transcript at the H16 locus which is polyadenylated. Concurrently, the expression of RNA exosome proteins is decreased, thereby preventing the normal degradation of such transcripts. The accumulation of these aberrant ncRNAs disrupts epigenetic patterns and leads to altered nucleolar morphology.

The increase in active transcription at the IGS was indicated by increased Pol II occupancy together with increased deposition of the H3K36me3 histone mark. Moreover, the marker of repressed chromatin, H3K9me3 was decreased at the IGS, suggesting that CDK12/13 inhibition results in a transcription-permissive environment that leads to the production of the aberrant ncRNAs along the IGS as well as disruption of nucleolar boundaries. Processed IGS transcripts play crucial roles in the establishment and maintenance of heterochromatin configuration at the promoters of rDNA arrays^24^. In the absence of CDK12/13 activity, heterochromatin configuration at rDNA is compromised, together with mislocalization of key nucleolar proteins. Thus, these CDKs may have protective roles in ensuring nucleolar organization by regulating Pol II transcription at the IGS.

We observed that the aberrant expression of IGS transcripts was mediated by Pol II, whereas Pol I remained confined to the 47S rDNA region. Low level transcription by Pol II at the IGS was present during steady state conditions; however, with CDK12/13 inhibition, Pol II occupancy increased, with elevated transcriptional activity and increased expression of IGS transcripts such as that at H16. The associated increase in Pol II initiation-related pS5 accompanied by a decrease in the elongation-related pS2 binding at H16 and at various other IGS loci such as H18 and H27 are consistent with effects at protein coding genes^7^, where perturbation of CDK12 is associated with an elongation defect with resultant accumulation of Pol II S5 at transcription start sites (TSS). Such an increase is manifested by the accumulation of nascent RNA reads both upstream and at the TSS. Moreover, as Pol II is less capable of elongation, it is gradually released from chromatin to increase the pool of free Pol II molecules that engage in transcription initiation.

We also noted through northern blot analysis and sequencing, the H16 ncRNA was produced in the forward orientation. The absence of transcripts between the end of 28S and the 3’ ETS plus the lack of rescue of the H16 ncRNA with Pol I inhibition suggest that the aberrantly increased H16 RNA is not readthrough from the 28S but arises from Pol II activity. CDK9 may also be involved in the same phenomenon, as we also observed sense transcripts emanating from the H16 region upon CDK9 inhibition. It is conceivable that CDK12/13 keeps the level of Pol II-mediated transcription at levels that ensure homeostasis. In protein-coding genes, Pol II transcribes RNA in both directions upon binding to the promoter, however, it only elongates productively into the gene body, while the transcript produced in the opposite direction is degraded^54^. It is possible that Pol II transcription follows the same pattern at IGS loci. The efforts of Pol II during steady states could be directed mainly towards generating the functional reverse ncRNA transcript barrier that prevents Pol I from producing sense intergenic ncRNAs through run-on transcription from the 28S rRNA genes^31^, while lower-level forward transcripts are likely targeted for degradation by the exosome. We postulate that during Pol II stalling, as occurs with the elongation defect, upstream antisense RNAs are outcompeted by forward transcripts and accumulate due to concomitant dysfunction of the degradation complexes. How and whether their aberrantly increased expression leads to CDK12/13-mediated cytotoxicity is currently unclear.

A key question is whether the observed effects on rRNA transcription were modulated by the three main targets of E9, CDK12, CDK13 or CDK9. The use of this inhibitor circumvented the compensatory effects that may have arisen from kinases with similar functions such as CDK12 and CDK9. CDK12 inhibition results in gene length-dependent elongation defects, at protein-coding genes, with premature cleavage at alternative polyadenylation sites and early termination of long (>45 kb) genes^7,52^. Indeed, members of the TRAMP complex, ZCCHC7 and TENT4A are 237 kb and 43 kb long respectively, while MTREX is 117 kb in length, so it is to be expected that their expression would be curtailed. Similarly, CDK13 has recently been shown to promote the degradation of aberrantly terminated protein-coding gene transcripts by phosphorylating a protein subunit of the PAXT complex^13^. Therefore, these kinases could be required for the prevention of aberrant transcript accumulation at the IGS region. Indeed, a CDK12 mutation at the covalent binding site that disrupts its interaction with E9 did not lead to IGS transcript accumulation and both kinases needed to be depleted to exhibit the effects of E9. Thus, both kinases are involved, CDK12 to protect Pol II from aberrant transcription, and CDK13 to degrade aberrant transcripts and direct them to the exosome. CDK9 inhibitors have been shown to induce a nucleolar stress phenotype by indirectly dissociating Pol I from rDNA though disruption of the Pol I-recruiting protein, TCOF^55^. We also observed nucleolar condensation and abnormalities of all three functional compartments, as well as contraction of 28S and 18S RNA staining with the E9 inhibitor. However, there was no change in the RNA expression of the rRNA genes, nor was occupancy of Pol I at rDNA genes compromised. While the effects seen here could be mediated solely by CDK9 inhibition, our findings with genetic and chemical degradation of CDK12 and CDK13 suggest that additional transcriptional CDK inhibitors may also play similar or concerted roles in regulating rDNA transcription and nucleolar integrity.

In summary, our findings suggest that Pol II regulation by CDKs 12 and 13 is not just essential for protein-coding genes but also in maintaining homeostasis of ribosomal RNA. CDK12/13 inhibition leads to profound consequences in the nucleolus, with significant increases in Pol II occupancy at the IGS region, accumulation of aberrantly polyadenylated transcripts and impairment of nucleolar activity. Moving forward, it will be essential to better understand the functional role of rDNA transcription in stress resolution and overall cellular homeostasis, particularly as transcriptional CDK inhibitors advance through clinical trials.

## Materials and Methods

### Cell Culture

Human OS cell lines (143B, U2OS, HOS and SaOS2) obtained from the American Type Culture Collection (ATCC) were grown in 1X DMEM (Thermo Fisher Scientific, 11995065) and supplemented with 10% FBS (Thermo Fisher Scientific, 10437028) and 1% penicillin/streptomycin (Thermo Fisher Scientific, 15140163). All cell lines were routinely tested for mycoplasma and authenticated through short tandem repeat analyses.

### Compounds

E9, THZ-531, BSJ-4-116, BSJ-4-23, BSJ-4-23NC, THZ1 and YKL-5-124 were synthesized and provided by Dr. Nathanael Gray (Dana-Farber Cancer Institute, Stanford University). dTAG13 (Cat #: 66-055) and Actinomycin D (Cat #: 11805017) were purchased from Fisher Scientific, CR8 (Cat #: C3249) and CX-5461 (Cat #: 5.09265) from Sigma Aldrich, NVP-2 (Cat #: HY-12214A) and RG-7834 (Cat #: HY117650A) from MedChem Express, and flavopiridol from Selleck Chemicals (Cat #: S2679).

### Cell transfection

For gene silencing, cells were transfected with small interfering RNA (siRNA) using Lipofectamine RNAiMAX (Thermo Fisher Scientific, 13778150) according to the manufacturer’s instructions. Briefly, cells were seeded at a density ensuring 50–70% confluency at the time of transfection. siRNAs were mixed with Lipofectamine RNAiMAX at a 1:2 ratio to achieve a final concentration of 10 nM. The mixture was incubated at room temperature for 10–15 minutes before being added dropwise to the cells. Six hours post-transfection, the medium was replaced with fresh complete growth medium, and cells were incubated for 48 hours at 37°C in 5% CO₂. Knockdown efficiency was assessed by RT-qPCR, using GAPDH as a loading control. The sequences of the siRNAs used in this study are listed in Supplemental Table 1.

### dTag knock-in

The original EGFP-2A-Puro cassette was removed from the pCRIS-PITChv2-FBL plasmid (Addgene, #63672) through restriction digestion using MluI-HF enzymes (New England Biolabs (NEB)). The BSD^R^-P2A-2xHA-FKBP12^F36V^-linker cassette, generated with flanking sequences homologous to the CDK12 locus, was assembled into the pCRIS-PITChv2 backbone using the NEBuilder HiFi DNA Assembly Cloning Kit (NEB), following the manufacturer’s protocol. To target CDK12, guide RNAs were inserted into the pX330A-1×2 plasmid (Addgene, #58766). Additionally, the PITCh-specific sgRNA sequence from pX330S-2-PITCh (Addgene, #63670) was cloned into pX330A-1×2-CDK12 using Golden Gate assembly (NEB). The resulting construct, pX330A-CDK12/PITCh, contained the PITCh sgRNA, the CDK12-targeting sgRNA, and Cas9. Next, 143B cells were seeded onto 10 cm plates and co-transfected with 1 µg of pCRIS-PITChv2-BSD^R^-dTAG-CDK12 and 2 µg of pX330A-CDK12/PITCh, using Lipofectamine 2000 (Thermo Fisher Scientific, 11668027) in accordance with the manufacturer’s instructions. Six hours post-transfection, fresh growth medium was added, and the cells incubated for an additional 48 hours following selection using 10 µg/mL of blasticidin to isolate individual clones into 96-well plates. Successful knockout of endogenous CDK12 and exogenous FKBP12F36V-CDK12 fusion protein expression were confirmed through Western blot analysis.

### Immunofluorescence microscopy

Cells were seeded on glass coverslips in 12-well plates at a seeding density of 5 × 10^4^ cells/well and treated with DMSO or the indicated compound. Following treatment, cells were washed in PBS and fixed in 4% formaldehyde in PBS for 15 min at room temperature (RT). Cells were permeabilized in 0.1% Triton X-100 in PBS for 10 min, washed and incubated in PBS containing 0.05% Tween-20 and 5% BSA (PBS-Tween-BSA) for at least 1 hour to block nonspecific binding. Cells were then incubated overnight at 4°C with primary antibody in PBS-Tween-BSA, extensively washed and incubated for at least 1 hour with fluorescently conjugated secondary antibody and counterstained with DAPI. Images were acquired on a Zeiss AXIO Imager Z1 fluorescence microscope using a 100x immersion objective, equipped with AxioVision software. Antibodies can be found in Supplemental Table 1.

### qRT-PCR

Total RNA was isolated from cells using the RNeasy Mini Kit (QIAGEN). For total RNA expression measurements, 1.5 μg of RNA was reverse transcribed using SuperScript IV VILO Master Mix (Life Technologies) according to the manufacturer’s protocol. For polyadenylated RNA expression quantification, 1.5 μg of RNA was reverse transcribed using SuperScript III Reverse Transcriptase (Life Technologies) according to manufacturer’s protocol and supplemented with 50 μM oligo(dT)_20_ primer. Quantitative PCR was carried out using the QuantiFast SYBR Green PCR kit (Qiagen) and analyzed on an Applied Biosystems StepOne Real-Time PCR System (Life Technologies). Each individual biological sample was qPCR-amplified in technical triplicate and biological duplicate or triplicate. Gene expression was normalized to GAPDH. Fold change in expression was calculated according to the ΔΔCt relative quantification method. Primer sequences were generated using the human genome alignment, hg38 and can be found in Supplemental Table 1.

### Northern blotting

Total RNA was denatured in 1x RNA loading buffer (2x, 98% w/v formamide with 10mM EDTA and 300 µg/µl bromophenol blue) by boiling at 90°C for 90 seconds, cooled on ice and resolved using a 1% agarose gel in 1x NorthernMax MOPS buffer (Thermo Fisher Scientific, AM8671) at 50V for 2 hours or until dye front migrated to ¾ down the gel. The gel was stained in 1x SYBR gold (Life Technologies S11494) to visualize the RNA, incubated in 10 mM NaOH for 20 min with shaking, rinsed with ddH2O, incubated with 20x UltraPure SSC (Life Technologies 15557044) for 5 minutes, and then with 10x SSC buffer for 40 minutes. The RNA was transferred onto a nylon (N+) membrane (GE Healthcare RPN203B) by diffusion blotting. The membrane was crosslinked and pre-hybridized in UltraHyb Hybridization Solution (Thermo Fisher Scientific, AM8670) at 42 °C for 1 hour, then incubated overnight with 50 pmol IGS-specific probes. The probes were synthesized with corresponding sequence DNA oligos and [γ-32P] UTP (Perkin Elmer) by using the MAXIscript T7 Transcription Kit (Invitrogen AM1312) according to the manufacturer’s protocol. Membranes were washed twice with 20 mL 6x SSC for 5 minutes at 42 °C, twice with 20 mL 2x SSC, and twice with 20 mL 1x SSC, wrapped in Saran wrap, exposed to a phosphor screen overnight, and visualized by phoshorimaging.

### RNA In-situ Hybridization

Cells were seeded on glass coverslips in 12-well plates at a seeding density of 5 × 10^4^cells/well, treated with DMSO or the indicated compound/siRNA (6 hours, 200nM E9; 72 hours, 20nM siRNA; 6 hours, 10 μM RG-7834). Following treatment, cells were washed in PBS and fixed in 4% paraformaldehyde for 10 minutes. Next, cells were permeabilized with 100% ice cold methanol for 10 minutes followed by 10 minutes in ice cold 70% ethanol, washed in 1M Tris-HCl, pH 7.5 and incubated overnight with probe hybridization buffer (2X SSC, 500 μg/ml yeast tRNA, 25% formamide, 0.005% BSA, 5% dextran sulfate, 1:5000 of 1 μg/μl fluorescently labeled probe). Slides were serially washed in 4X SSC, 2X SSC, and 2X SSC and mounted using fluoromount solution containing DAPI. Probes sequences can be found in Supplemental Table 1.

### Western Blotting

Cells were collected using 18 cm Cell Lifters (GeneMate) and lysed by boiling in 4% SDS buffer. Protein lysates were diluted to 1.33% SDS with water and concentrations determined using the Biorad DC Protein Assay kit (BioRad). Whole cell lysates were resolved on 4-12% BisTris gels (Invitrogen) and transferred to nitrocellulose membranes (BioRad). Following transfer, membranes were blocked in 5% milk dissolved in Tris-buffered saline containing 0.2% Tween-20 (Boston BioProducts). Antibody information can be found in Supplemental Table 1. Target proteins were detected by chemiluminescence using GeneMate Blue Ultra Autoradiography film (VWR).

### Chromatin Immunoprecipitation-quantitative PCR (ChIP-qPCR)

HOS or SaOS2 cells (1 x 10^7^) were crosslinked for 10 minutes at room temperature with 1% formaldehyde (Thermo Fisher Scientific, 28908) in PBS followed by quenching with 0.125 M glycine for 5 minutes, washed twice in PBS, and cell pellets flash frozen and stored at −80 °C. Forty μl of protein G Dynabeads per sample (Invitrogen) were blocked with 0.02% Tween20 (w/v) in PBS, loaded with antibody (2 μg for Pol I, and 5 μg for Pol II, pS5-Pol II, pS2-Pol II, H3K9me3 and H3K36me3) and incubated overnight at 4 °C. Crosslinked cells were lysed, placed in sonication buffer with 0.2% SDS, placed on ice and chromatin was sheared using a Misonix 3000 sonicator (Misonix) at the following settings: 10 cycles, each for 30 seconds on, followed by 1 minute off, at a power of approximately 20 W. The lysates were centrifuged for 10 minutes at 4 °C, supernatants collected and diluted with an equal amount of sonication buffer without SDS. The sonicated lysates were incubated overnight at 4 °C with the antibody-bound magnetic beads, washed with low-salt buffer (50 mM HEPES-KOH (pH 7.5), 0.1% SDS, 1% Triton X-100, 0.1% sodium deoxycholate, 1 mM EGTA, 1 mM EDTA, 140 mM NaCl and 1× complete protease inhibitor), high-salt buffer (50 mM HEPES-KOH (pH 7.5), 0.1% SDS, 1% Triton X-100, 0.1% sodium deoxycholate, 1 mM EGTA, 1 mM EDTA, 500 mM NaCl and 1× complete protease inhibitor), LiCl buffer (20 mM Tris-HCl (pH 8), 0.5% NP-40, 0.5% sodium deoxycholate, 1 mM EDTA, 250 mM LiCl and 1× complete protease inhibitor) and Tris-EDTA buffer. DNA was then eluted in elution buffer (50 mM Tris-HCl (pH 8.0), 10 mM EDTA, 1% SDS), and high-speed centrifugation was performed to pellet the magnetic beads and collect the supernatants. The crosslinking was reversed overnight at 65 °C. RNA and protein were digested using RNase A and proteinase K, respectively, and DNA was purified with phenol chloroform extraction and ethanol precipitation. qPCR was carried out using the QuantiFast SYBR Green PCR kit (Qiagen) and analyzed on an Applied Biosystems StepOne Real-Time PCR System (Life Technologies). Each individual biological sample was qPCR-amplified in technical duplicate and biological duplicate. Chromatin enrichment was calculated as previously described using (percentage of input/IgG) = (percentage of input for protein immunoprecipitation)/(percentage of input for mock IgG immunoprecipitation)^31^.

### Nascent RNA quantification

Cells were exposed to DMSO or E9 at 200 nM for 6 hours and nascent RNA was labeled using 4-thiouridine (4sU) incorporation and purified as described by Schwalb *et al*. (2016)^56^. Briefly, cells were incubated with 500 µM 4sU for 10 minutes to metabolically label newly synthesized RNA. After labeling, total RNA was extracted using the RNeasy Mini Kit (Qiagen) and treated with DNase I to eliminate genomic DNA contamination. For purification of nascent RNA, 4sU-labeled transcripts were selectively biotinylated by incubating total RNA with biotin-HPDP (Pierce, 21341) in 10 mM Tris-HCl (pH 7.4), 1 mM EDTA, and 0.2 mg/mL biotin-HPDP in dimethylformamide (DMF) for 1 hour at room temperature in the dark. Unreacted biotin-HPDP was removed using chloroform-phenol extraction, and labeled RNA was captured on streptavidin-coated magnetic beads (Dynabeads MyOne Streptavidin C1, Thermo Fisher Scientific) for 30 minutes at room temperature with gentle rotation. Beads were then washed extensively with high-salt buffer (100 mM Tris-HCl pH 7.4, 10 mM EDTA, 1 M NaCl, 0.1% Tween-20) to remove unlabeled RNA. Biotinylated nascent RNA was eluted from the beads using 100 mM DTT and purified with RNeasy MinElute columns (Qiagen). Purified RNA was then used for RT-qPCR.

### Polyadenylated Tail Length Assay

To detect the polyadenylated tail of the H16 transcript, we utilized the USB Poly(A) Tail-Length Assay Kit (Life Technologies) per the manufacturer’s protocol. Primer sequences can be found in Supplemental Table 1. PCR reactions were resolved on a 2% agarose gel. Lane 5 (PCR amplified DNA using H16 Poly(A) primer in E9-treated cells) was excised from the gel, DNA was purified using the Nucleospin Gel and PCR Clean Up Kit (Takara Bio), and purified DNA submitted to Genewiz for standard Sanger sequencing.

### RNA immunoprecipitation-quantitative PCR (RIP-PCR)

HOS cells (1 x 10^7^) were disrupted by using a lysis buffer composed of 20 mM HEPES pH 7.9, 150 mM NaCl, 1 mM MgCl2, 1 mM EGTA, 10% glycerol, 1% Triton-X100, 0.1% sodium deoxycholate, in the presence of a complete protease inhibitor cocktail devoid of EDTA (Roche) and 20 U/ml RNase inhibitor (Invitrogen) for 45 minutes, sonicated briefly and then centrifuged at 15,000 g for 15 minutes at 4°C. The supernatants were subjected to pre-clearing using protein G beads for 30 minutes, followed by incubation with anti-MTREX, anti-EXOSC2 or control IgG-coated beads for 4 hours at 4°C. The beads were washed four times with lysis buffer and then resuspended in 100 μl of NT2 buffer (50 mM Tris–HCl pH 7.4, 150 mM NaCl, 1 mM MgCl2, 0.05% NP-40) containing 10 U of DNase and incubated at 37°C for 10 minutes. The RNA-protein complexes were eluted using elution buffer (10 mM Tris–HCl pH 8.0, 1 mM EDTA, 1% SDS, 20 U/ml RNase inhibitor) at 65°C for 10 minutes and the eluates then treated with Proteinase K (1 mg/ml) at 50°C for 5 hours. RNA was isolated using Trizol reagent (Invitrogen) and analyzed using qRT-PCR.

### Cell Viability Assay

For TENT4A knockdown, 20nM siRNA was complexed with RNAiMAX according to manufacturer’s protocol. 5 x 10^3^ cells/well were added to the complexed siRNA and allowed to grow for 72 hours. For RG-7834 viability assay, cells were plated in 96-well plates at a seeding density of 5 x 10^3^ cells/well. After 20 hours, cells were treated with 10 μM RG-7834. 72 hours after drug administration cells were analyzed using the CellTiter-Glo Luminescent Cell Viability Assay (Promega) according to the manufacturer’s instructions. Cell viability assays were performed in biological and technical triplicates. Cell viability was normalized to 72-hour treatment with the DMSO solvent.

### Analysis of RNA-sequencing datasets

Publicly available RNA sequencing data was downloaded from Sequence Read Archive (SRA) in fastq format as mentioned in Supplemental Table 2 The data was processed using the framework for analyzing rDNA sequence as previously described^57^. Briefly, a custom build of genome assembly was created by concatenating a full, non-repeat masked human rDNA repeat (GenBank accession no. U13369) with human genome GRCh38 and External RNA Controls Consortium (ERCC) spike-in sequences. The RNA sequences were aligned to this custom genome with STAR (v2.7)^58^. Spike-in normalization factor was calculated as 100000/number of reads mapped to the ERCC sequences for each sample. The aligned sequences were then used to generate strand-specific bigWig tracks using the bamCoverage function in deepTools^59^ with the --filterRNAstrand parameter and scaled with spike-in normalization factor. Integrative Genomics Viewer (IGV)^60^ was utilized to visualize the bigWig tracks. Differential expression information for genes of interest were obtained from the microarray signal intensities or FPKM values as described in the corresponding publication of source data^7,15^ and the fold change heatmap was generated with GraphPad Prism. Average gene lengths for the RNA exosome, NEXT, PAXT and TRAMP component genes were calculated as mean of gene lengths in KB, obtained from GRCh38 annotation, and visualized as a violin plot.

### Statistical Analysis

Each experiment was performed in biological duplicates or triplicates. Comparison between two groups was analyzed using the two-tailed unpaired Student’s t-test. Comparison of three or more groups was analyzed using the one-way ANOVA followed by Dunnett’s multiple comparisons test. Results are shown as mean ± SEM. GraphPad Prism 4.0 software was used for data plotting.

## Data Availability

Supplementary table 1 includes all nucleotide sequences used in this work. Further information and requests for resources and reagents should be directed to and will be fulfilled by the Lead Contact, Rani E. George.

## Acknowledgements

*Author contributions*: Conceptualization (RBS; MH; REG), Investigation (RBS; MV; MH; CC; BS), Formal Analysis (RBS; MH; UB; SD; GEZ; PCS), Funding acquisition (REG; NSG), Resources (REG; TZ; NSG; SB), Writing – original draft (RBS; MH; REG), Writing – review & editing (RBS; UB; REG), Project administration (REG).

---

**Supplemental Figure 1.**
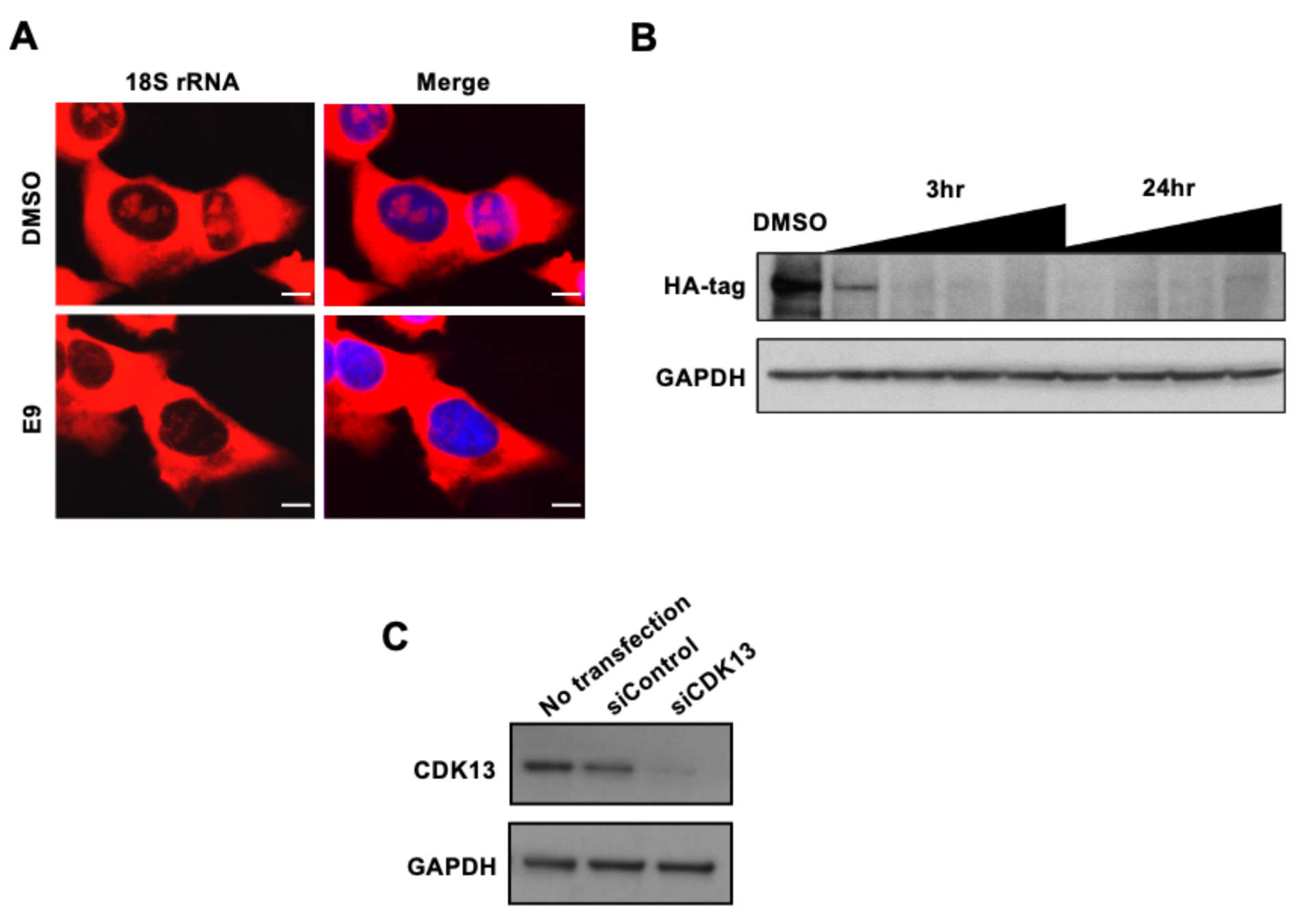
CDK12/13 inhibition causes nucleolar contraction. (A) RNA-ISH analysis of 18S rRNA in OS cells treated with E9 (200 nM x 6 h). Scale bar, 10 μm. (B) Western blot (WB) analysis of CDK12 protein expression in 143B CDK12 dTAG cells treated with DMSO or 1 µM dTAG13 for 3 or 24 h. HA antibody was used for detection of HA-CDK12. (C) WB analysis of CDK13 protein expression in 143B CDK12 dTAG cells transfected with siControl or siCDK13.

**Supplemental Figure 2.**
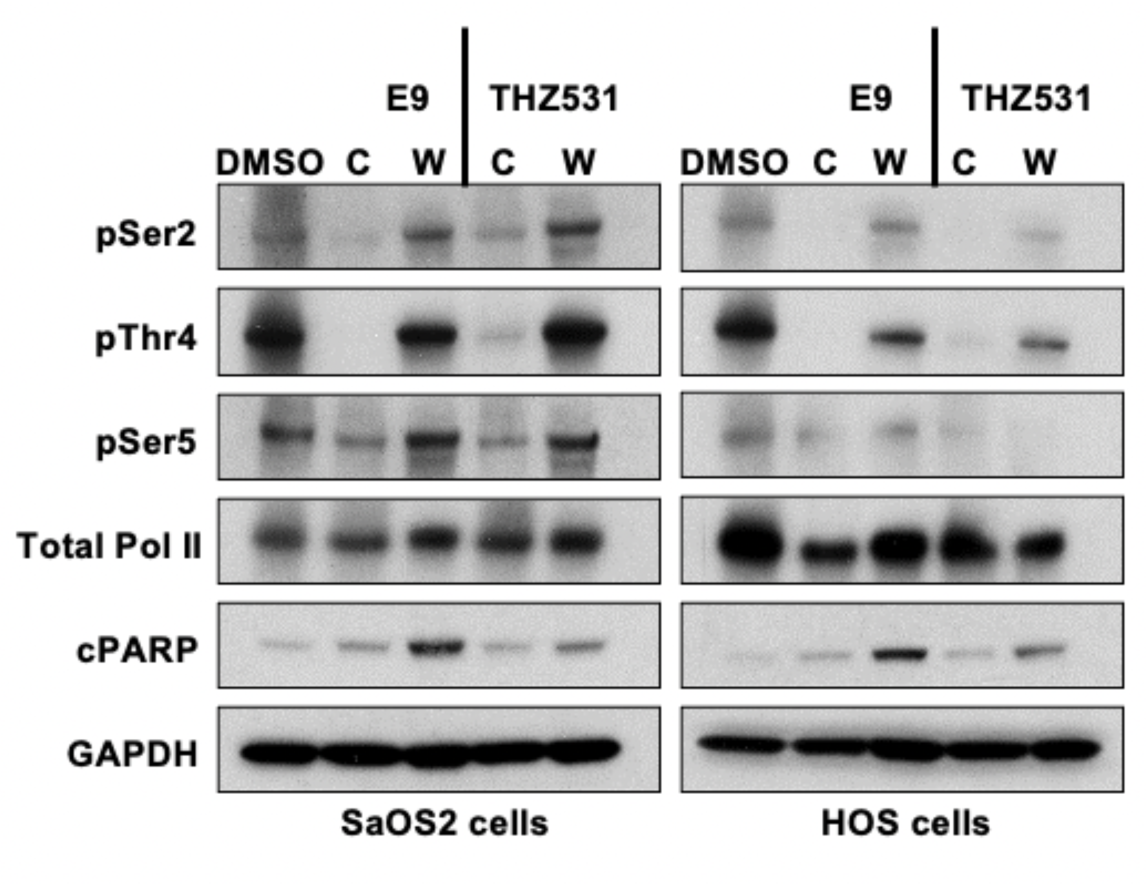
CDK12/13 inhibition affects Pol II phosphorylation. WB analysis of Pol II phosphorylation and cleaved PARP in SaOS2 and HOS OS cells treated with E9 (200 nM x 6 h) or THZ531 (400 nM x 6 h). C: continuous treatment for 6 h followed by sample collection; W: drug washout after 6 h treatment followed by sample collection after 24 h. GAPDH was used as loading control.

**Supplemental Figure 3.**
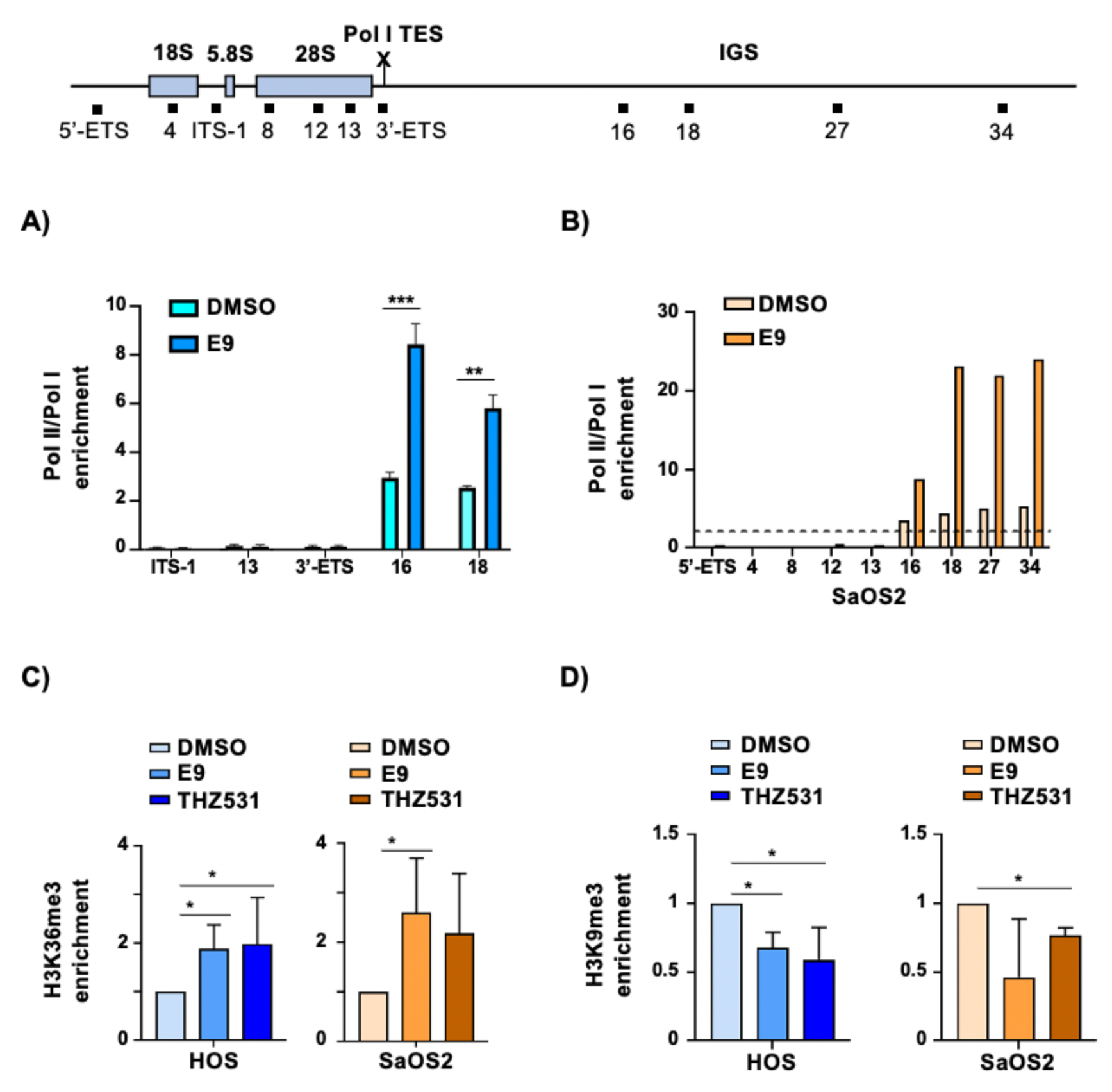
CDK12/13 inhibition creates a transcriptionally permissive chromatin environment within the IGS enabling Pol II activity. (A) ChIP-qPCR analysis of Pol II and Pol I occupancies at the 28S (H13), ITS-1, 3’ETS, H16 and H18 regions in HOS cells treated with E9 (200 nM x 6 h). The ratio of Pol II to Pol I binding was calculated as (% of input for Pol-II immunoprecipitation)/(% of input for Pol-I immunoprecipitation)/(% of input for mock IgG immunoprecipitation). Data represent mean ± S.D. of 2 independent replicates. \*\**P* < 0.001, and \*\*\**P* < 0.0001, Student’s t test. (B) ChIP-qPCR analysis of Pol II and Pol I occupancies at the 18S (H4), 28S (H8, H12 and H13) and IGS regions in SaOS2 cells treated with E9 (200 nM x 6 h). Data represent mean ± S.D. of 2 independent replicates. Pol-II/Pol-I ratio calculated as in (A). (C-D) ChIP-qPCR analysis of H3K36me3 (C) or H3K9me3 (D) occupancies at the H16 region in HOS or SaOS2 cells treated with E9 (200 nM x 6 h) or THZ531 (500 nM x 6 h). Data represent mean ± S.D. of 2 independent replicates. \**P* < 0.05, one-way ANOVA followed by Dunnett’s multiple comparisons test.

**Supplemental Figure 4.**
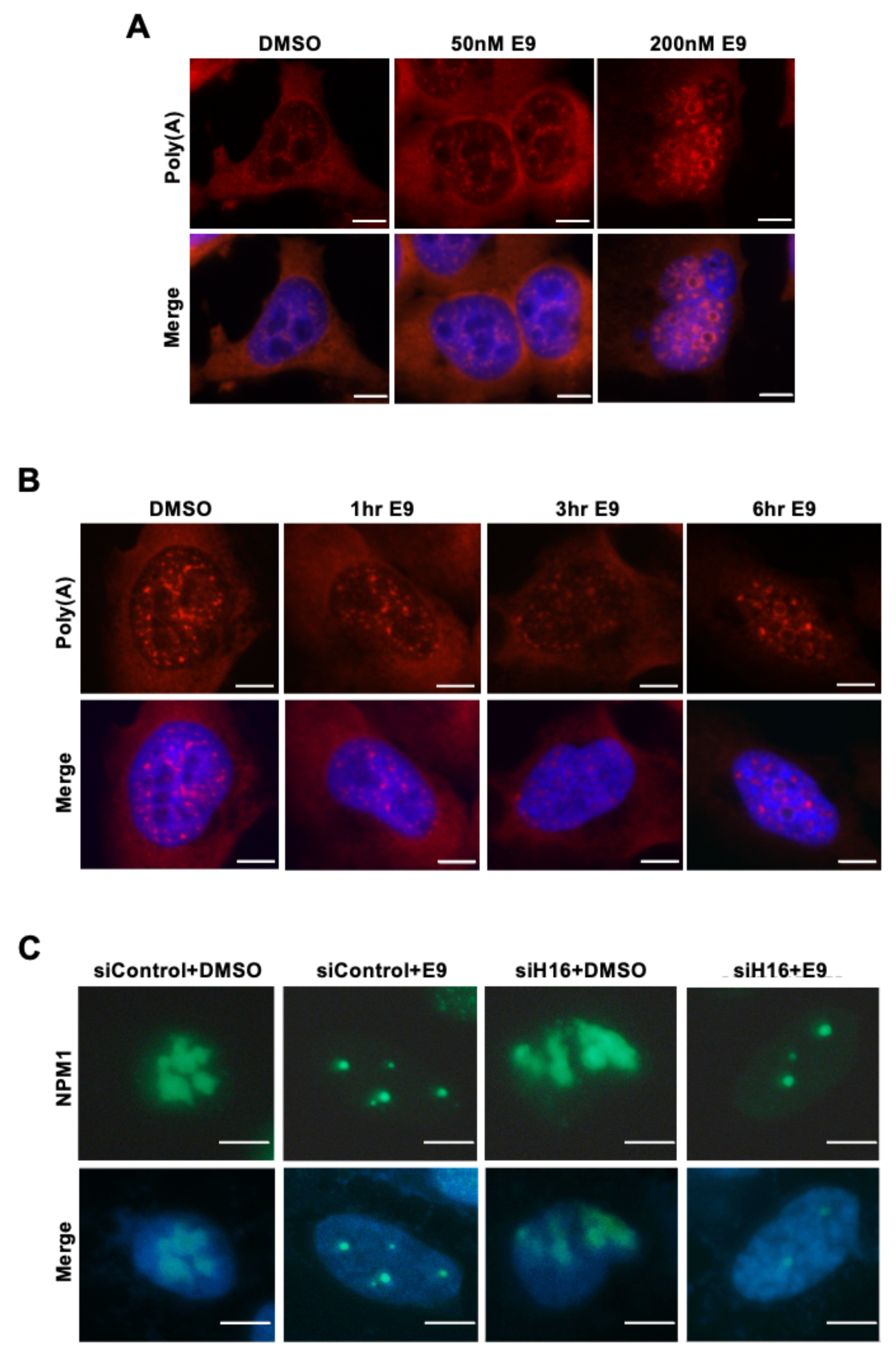
pAR rings increase in an E9 time- and concentration-dependent manner. (A) IF analysis of poly(A) in OS cells incubated with E9 (50 nM or 200 nM x 6 h). Scale bar, 5 μm. (B) IF analysis of poly(A) in OS cells incubated with E9 (200 nM x 1, 3 or 6 h). Scale bar, 5 μm. (C) IF analysis of NPM1 in OS cells transfected with siRNA against H16 RNA and incubated with E9 (200 nM x 6 h). Scale bar, 5 μm.

**Supplemental Figure 5.**
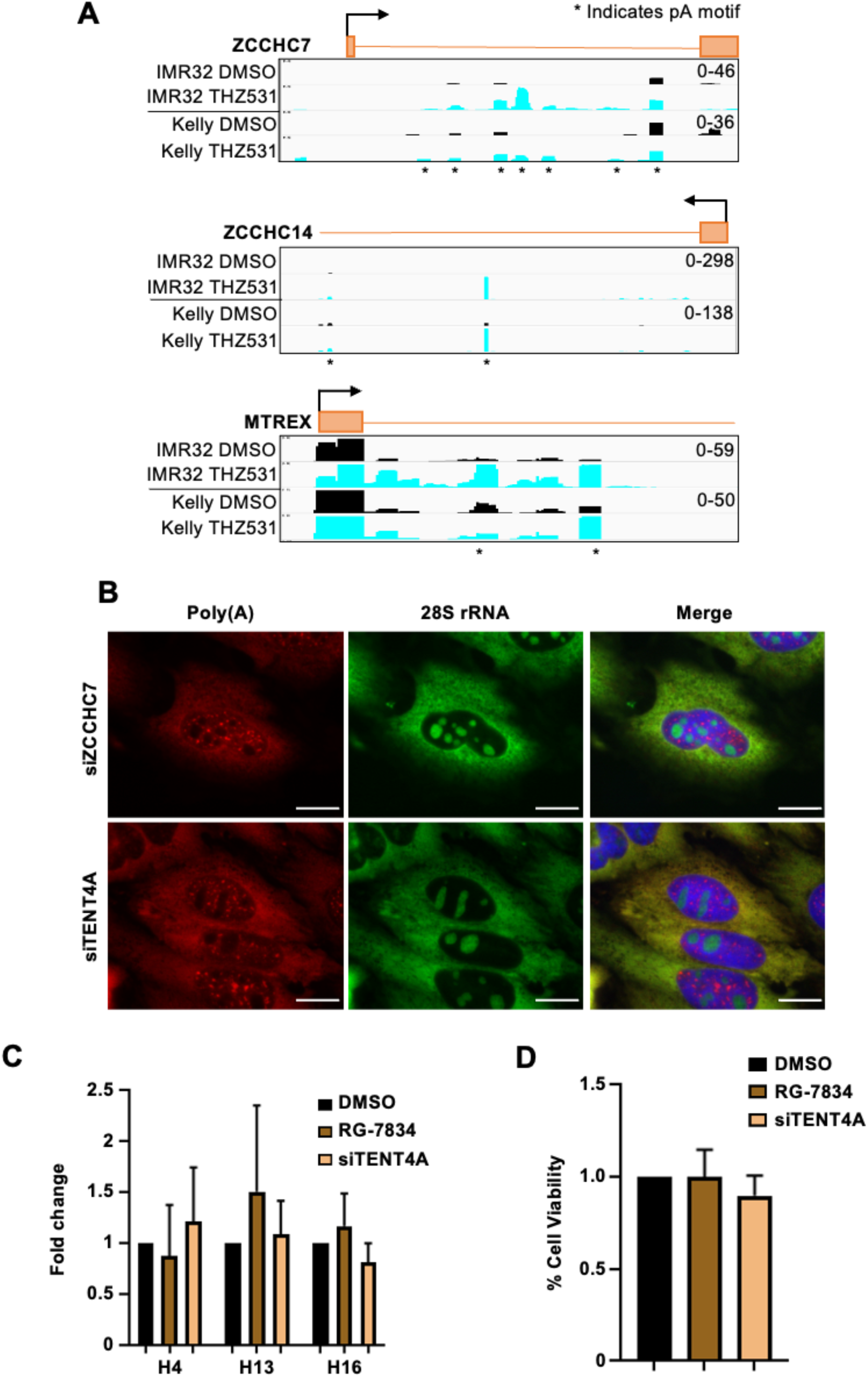
ZCCHC7 or TENT4A inhibition do not affect rRNA expression and cell viability. (A) 3’-Poly(A) RNA-sequencing tracks of ZCCHC14, MTREX and ZCCHC7 genes in IMR32 and Kelly cells treated with THZ531 (400 nM x 6 h) and DMSO. Intronic polyadenylation sites are marked with *. (B) IF analysis of poly(A) and 28S rRNA in OS cells transfected with siRNA against ZCCHC7 or TENT4A mRNA. Scale bar, 5 μm. (C) RT-qPCR analysis of 18S (H4), 28S (H13) and H16 region in OS cells treated with RG-7834 or transfected with siRNA against TENT4A mRNA. (D) Cell viability in OS cells treated or transfected as in (C). Data represent mean ± S.D. of 3 independent replicates.

