## Supplemental table 1 for "CDK12/13 inhibition disrupts nucleolar morphology and promotes aberrant expression of IGS transcripts"

Supplemental Table 1: Oligonucleotide sequences, FISH and Northern Blot probes, and antibodies used in this study.

| Oligo Name | Sequence - 5’ to 3’ | Length |
|---|---|---|
| 5'-ETS_F | GGCGGTTTGAGTGAGACGAGA | 21 |
| 5'-ETS_R | ACGTGCGCTCACCGAGAGCAG | 21 |
| ITS-1_F | CGAGAGCCGAGAACTCGGGAG | 21 |
| ITS-1_R | CGCACACCAACGACACGCCCT | 21 |
| 3'-ETS_F | GTTCGGCGCCGTCCGTCCTT | 20 |
| 3'-ETS_R | GAGGCGGGAACCGAAGAAGC | 20 |
| rRNA04_F | CGACGACCCATTCGAACGTCT | 21 |
| rRNA04_R | CTCTCCGGAATCGAACCCTGA | 21 |
| rRNA08_F | AGTCGGGTTGCTTGGGAATGC | 21 |
| rRNA08_R | CCCTTACGGTACTTGTTGACT | 21 |
| rRNA12_F | GAGCTCAGGGAGGACAGAAA | 20 |
| rRNA12_R | AGGTCAGAAGGATCGTGAGG | 20 |
| rRNA13_F | CACCGCACGTTCGTGGGGAA | 20 |
| rRNA13_R | ACAAACCCTTGTGTCGAGGGC | 21 |
| IGS13.4_F | ACCGACGAACCGCGGGTGGC | 20 |
| IGS13.4_R | CGGAGGCCCGACCGAGGAGA | 20 |
| IGS14.3_F | CGGCTCTTGAGACTTAGCCGCTG | 23 |
| IGS14.3_R | AGGTGCCGACCGAGACGGGGA | 21 |
| IGS15_F | GTCGACCGGCGGGCCTTCT | 19 |
| IGS15_R | TCGACCCCGGAGGTGCCGA | 19 |
| IGS16_F | ACACACACACACCCCGTAGT | 20 |
| IGS16_R | GAAATGGGGCTTCGATACAT | 20 |
| IGS18_F | GTTGACGTACAGGGTGGACTG | 21 |
| IGS18_R | GGAAGTTGTCTTCACGCCTGA | 21 |
| IGS20_F | GTAGCCTTGGGCTTCTCTCC | 20 |
| IGS20_R | AGTTTTCAGCCCCAACACAC | 20 |
| IGS22_F | CAGTGGCTCACGTCTGTCAT | 20 |
| IGS22_R | CGCCTGACTCCATTTCGTAT | 20 |
| IGS24_F | CCCGCGCACATAATAACTAA | 20 |
| IGS24_R | AAATCACTCCTCACGGGAAC | 20 |
| IGS27_F | CCTTCCACGAGAGTGAGAAG | 20 |
| IGS27_R | GACCTCCCGAAATCGTACAC | 20 |
| IGS30_F | GGTCTCTGCGTCTCGCTATC | 20 |
| IGS30_R | TGAAGAATTCAGGCCTTGGT | 20 |
| IGS32_F | AAAAGCTGGCCGATCTGAAT | 20 |
| IGS32_R | CGTCTGTTCAGCTATTTTGCAG | 22 |
| IGS34_F | CCATGCCTTCGACTCTGTAA | 20 |
| IGS34_R | GTACTGTGCCAAATCGGAAA | 20 |
| IGS36_F | TCCACTCCCAAGTTCAGTGG | 20 |
| IGS36_R | CGAGGGAACCCAAGGTAGAG | 20 |
| IGS38_F | CTCACAGAGGAAGGGAGCAC | 20 |
| IGS38_R | AACAAGGGAGGGAGGAACTT | 20 |
| IGS40_F | TTCTCCTTGGTCAGGGGTTT | 20 |
| IGS40_R | CAGGAAAGTCCCCAACAACA | 20 |
| **FISH probes** | **Sequence - 5’ to 3’** | **Length** |
| 18S rRNA | TGACTCTAGATAACCTCGGGCCGATCGCACG | 31 |
| 28S rRNA | TGGGAATGCAGCCCAAAGCGGGTGGTAAAC | 30 |
| **Northern blot probes** | **Sequence - 5’ to 3’** | **Length** |
| Forward | TAATACGACTCACTATAGACCGAGCGGCGTGTAAGAGTGCCCGTCGGGACGAGCCGGACCCGCCGCGTCCCCGTCTCGG | 79 |
| Reverse | TAATACGACTCACTATAGCCGAGACGGGGACGCGGCGGGTCCGGCTCGTCCCGACGGGCACTCTTACACGCCGCTCGGT | 79 |
| **Antibodies** | **Catalog number** | **Company** |
| NOLC1 | SC-374033 | Santa Cruz |
| FBL | #2639S | Cell Signaling |
| NPM1 | sc-271737 | Santa Cruz |
| H3K9me3 | AB8898 | Abcam |
| HA-tag | #2367S | Cell Signaling |
| GAPDH | #2118 | Cell Signaling |
| CDK13 | ABE1860 | Sigma Aldrich |
| pSer2 RNAPII | ab238146 | Abcam |
| pThr4 RNAPII | 61361 | Active Motif |
| pSer5 RNAPII | ab5131 | Abcam |
| Total RNAPII | ab26721 | Abcam |
| cPARP | #5625S | Cell Signaling |
| RNAPI | #24799S | Cell Signaling |
| H3K36me3 | ab9050 | Abcam |
| MTREX | A300615A | Thermo Fisher |
| EXOSC2 | ab181211 | Abcam |
