## Supplemental table 2 for "CDK12/13 inhibition disrupts nucleolar morphology and promotes aberrant expression of IGS transcripts"

Supplemental Table 2: BigWig files for figure 2D.

| File | GSE ID | Paper Ref |
|---|---|---|
| SRR10582276_HEK293T_DMSO_forward.bigwig | GSE141377 | PMID: 32917631 |
| SRR10582280_HEK293T_THZ531_forward.bigwig | GSE141377 | PMID: 32917631 |
| SRR10582283_THP_1_cells_DMSO_forward.bigwig | GSE141377 | PMID: 32917631 |
| SRR10582284_THP_1_cells_THZ531_forward.bigwig | GSE141377 | PMID: 32917631 |
| SRR10744478_KBM7_DMSO_sorted.spikenorm.forward.bw | GSE142405 | PMID: 32747809 |
| SRR10744493_KBM7_THZ531_sorted.spikenorm.forward.bw | GSE142405 | PMID: 32747809 |
| SRR13075283_Jurkat_cells_BSJ-4-23_forward.bigwig | GSE161650 | PMID: 33753926 |
| SRR13075285_Jurkat_cells_BSJ-4-23NC_forward.bigwig | GSE161650 | PMID: 33753926 |
| SRR13075289_Jurkat_cells_DMSO_forward.bigwig | GSE161650 | PMID: 33753926 |
| SRR13075290_Jurkat_cells_THZ531_forward.bigwig | GSE161650 | PMID: 33753926 |
| SRR8174597_Kelly_DMSO_forward.bigwig | GSE113314 | PMID: 30988284 |
| SRR8174598_Kelly_THZ531_6h_forward.bigwig | GSE113314 | PMID: 30988284 |
| SRR8174601_Kelly_E9R_DMSO_forward.bigwig | GSE113314 | PMID: 30988284 |
| SRR8174602_Kelly_E9R_THZ531_6h_forward.bigwig | GSE113314 | PMID: 30988284 |
| SRR9204213_143B_DMSO_sorted.spikenorm.forward.bw | GSE132233 | PMID: 31498151 |
| SRR9204214_143B_E9_sorted.spikenorm.forward.bw | GSE132233 | PMID: 31498151 |
